# REHE: Fast Variance Components Estimation for Linear Mixed Models

**DOI:** 10.1101/2021.02.03.429643

**Authors:** Kun Yue, Jing Ma, Timothy Thornton, Ali Shojaie

## Abstract

Linear mixed models are widely used in ecological and biological applications, especially in genetic studies. Reliable estimation of variance components is crucial for using linear mixed models. However, standard methods, such as the restricted maximum likelihood (REML), are computationally inefficient and may be unstable with small samples. Other commonly used methods, such as the Haseman-Elston (HE) regression, may yield negative estimates of variances. Utilizing regularized estimation strategies, we propose the restricted Haseman-Elston (REHE) regression and REHE with resampling (reREHE) estimators, along with an inference framework for REHE, as fast and robust alternatives that provide non-negative estimates with comparable accuracy to REML. The merits of REHE are illustrated using real data and benchmark simulation studies.

## 1 Introduction

The linear mixed model is a convenient and powerful tool for analyzing correlated data, with a wide range of applications in scientific research. It is especially useful for genetic studies of complex traits, including heritability estimation [29], genome-wide association studies (GWAS) [1], and network-based pathway enrichment analysis (NetGSA) [28]. Variance components estimation is an essential step when applying linear mixed models and the restricted maximum likelihood (REML) approach is the gold-standard for this task [23]. REML works by iteratively maximizing the residual likelihood with respect to the variance component parameters. During each iteration, REML computes the inverse of two *n* × *n* matrices, where *n* is the sample size of the data set. As a result, the computation for REML quickly becomes prohibitive for large sample sizes, especially when the correlations among observations are non-sparse – such as between-subject correlations from genetic relationships [16]. Despite efforts such as average information REML [8], Monte Carlo REML [21], and REML based on grid search [15] to improve its computational efficiency, REML is still not scalable to large data sets in many applications, particularly in genetic studies. This impedes the ability to estimate heritability for the HSHC/SOL data set since it has over ten thousand individuals and genetic correlations are mostly non-zero among individuals. On the other hand, consistency and asymptotic normality of REML estimates are large sample properties. As will be illustrated with simulation studies in Section 4 and in Supplementary Note 2.3, REML may be numerically unstable or may provide unreliable estimates. Given these shortcomings, fast and reliable estimators for linear mixed model variance components are desired.

When computational efficiency is a primary concern, moment estimators have been used frequently as alternatives to REML. These include analysis of variance (ANOVA), minimum norm quadratic unbiased estimation (MINQUE), and the Haseman-Elston (HE) regression estimator [26, 25, 14, 29]. These methods bypass the most time-consuming step in REML — the inversion of *n* × *n* matrices. Their key idea is to set up estimating equations by equating the mean squared errors to its expectation, the error variance. ANOVA, originated from ideas by R. A. Fisher in 1920s, has been well-established for estimating variance components. The resulting estimators are minimum variance quadratic unbiased [12], and minimum variance unbiased under normality assumptions on the random effects and the errors [11, 13]. MINQUE [25], which can be viewed as an extension of the ANOVA method, is equivalent to the first iteration of REML [27]. It relaxes the assumption of normality using estimating equations that rely on initial values for the variance components. The HE estimator, first introduced in [14], has been recently used for linear mixed model variance component estimations in genetic studies [29, 33]. Its simple idea and fast computation make it favorable when working with large and densely correlated data sets. A key limitation of these moment estimators is that they do not guarantee non-negative estimates for the variance components. This leads to difficulties in interpretation and downstream analyses.

To address the shortcomings of existing approaches in linear mixed model variance component estimation, we propose a new estimation method based on restricted Haseman-Elston (REHE) regression. REHE is computationally efficient, and ensures non-negative estimates of variance components that are comparably accurate to REML estimates. To accommodate the need for a strictly positive variance estimates in some applications, we also propose REHE with resampling (reREHE), which provides a positive estimate with high probability. Furthermore, to facilitate inference, we propose bootstrap confidence intervals for REHE estimates; our numerical experiments show that these confidence intervals are more robust than their REML counterparts. Finally, to further speed up the computation for inference, we also take advantage of correlation matrix sparsification [9] in the variance component estimation context. We demonstrate the application of proposed methods in three different contexts: heritability estimation, GWAS and NetGSA. We benchmark the proposed methods’ performance with simulation studies, and illustrate their advantages with data from the Hispanic Community Health Study/Study of Latinos (HCHS/SOL) [30, 4] and The Cancer Genome Atlas (TCGA) breast cancer data set [31].

The rest of the paper is organized as follows. In Section 2, we introduce the REHE estimator, discuss its properties and propose a bootstrap inference framework. We also introduce the reREHE estimator as an alternative for the REHE estimator. We demonstrate the performance of the REHE and reREHE estimators with real data applications in Section 3. In Section 4, we benchmark their performance with extensive simulation studies. Section 5 concludes the paper with discussions on the results and potential improvements. Additional results on REHE, reREHE and matrix sparsification are in the Supplementary Materials.

## 2 Methods

Consider a generic linear mixed model for an outcome vector *Y* of length *n*:

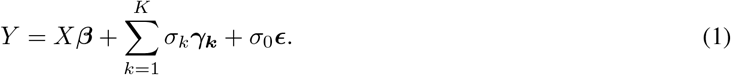

Here, *X* is an *n* × *p* design matrix for *p* covariates, and *β* is a *p*-dimensional fixed effect coefficient vector. For *k* = 1, 2, …, *K, γ*_*k*_ is a length *n* vector of random effects following *N*_*n*_(0, *D*_*k*_), where each *n × n* matrix *D*_*k*_ defines one source of relatedness among observations, and is assumed to be known. The noise *ϵ* is a length *n* vector following 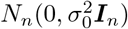. The parameters *σ*_*k*_’s (*k* = 0, 1, …, *K*) are the variance components. For 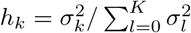 estimates the proportion of variation explained by *D*_*k*_. In the context of genetic studies, *D*_1_ is often the kinship matrix and *h*_1_ is referred to as heritability.

Our main objective is to estimate the variance components 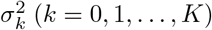. For expositional clarity, we assume the model has no fixed effect, and the outcome vector *Y* is centered. We also assume *K* = 1 such that *γ*_1_ explains all relatedness among observations. However, our methods can be easily extended to models with fixed effects (Section 2.2.3), or with more than one random effects. Denoting *D*_0_ = ***I***_*n*_ and *γ*_**0**_ = *ϵ*, model (1) becomes

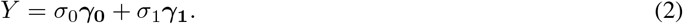

### 2.1 The Haseman-Elston Regression

The Haseman-Elston (HE) regression approach [14, 29, 33] estimates the variance components via the method of moments. Specifically, since model (2) implies that 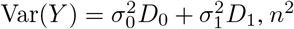 estimating equations are constructed:

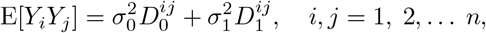

where 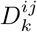 denotes the (*i, j*) entry of matrix *D*_*k*_ (*k* = 0, 1). Estimation of the variance components can thus be recast as a linear regression problem. Let 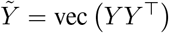 denote the vectorization of the *n* × *n* matrix *Y Y*^⊤^ by stacking its columns, 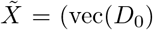 and 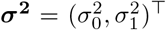. We then have 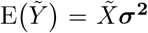. HE solves for variance components ***σ***^**2**^ by linear regression, specifically, by minimizing the residual sum of squares

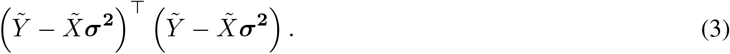

The resulting estimator has a closed form expression 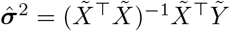.

The computational complexity of HE is *O*(*Kn*^2^) compared to *O*(*n*^3^) for REML. When the sample size *n* is large and the number of variance components *K* is small, as is typically the case in practice, HE offers substantial improvement in computation over REML. However, the ordinary least squares solution for variance components by HE is not guaranteed to be non-negative, leading to difficulties in downstream analyses and interpretation. In practice, negative estimates from HE are often truncated at zero; yet such naive truncation does not minimize the residual sum of squares in (3) within the parameter space 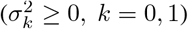. In addition, as will be illustrated in simulation studies in Section 4 and the Supplementary Note 2.3, naive truncation-based HE estimates generally has larger mean square error than estimates by REML and our proposed method, discussed next.

### 2.2 The Restricted Haseman-Elston Regression

To prevent negative estimation of variance components by HE, while preserving its computational efficiency, we propose a new variance components estimation method, termed the restricted Haseman-Elston (REHE) regression. Similar to HE, REHE is a moment estimator which regresses the empirical covariance of the observations on pre-specified correlation matrices that encode sample relatedness. However, instead of the ordinary least squares estimate by HE, REHE finds the non-negative minimizer of the residual sum of squares, ensuring sensible estimation of variance components (Supplementary Figure S10). Following (3), the REHE estimates of the variance components are expressed as:

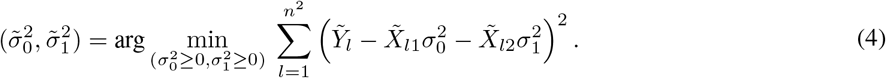

There is a closed form solution to (4) with only two variance components (Supplementary Note 1.1). With more than two variance components, iterative algorithms for non-negative least squares (NNLS) can easily solve (4) [18, 10, 17, 7]. The convexity of (4) guarantees the numerical solutions of different solvers converge to the same global minimizer. Using the R package *quadprog* (*v1*.*5-7*, 2), REHE estimation has approximately the same computational cost as HE, and is thus substantially faster than REML.

#### 2.2.1 Consistency and Asymptotic Normality

The REHE estimator is consistent and asymptotically normal under mild conditions. For simplicity, we assume *D*_1_ is sparse and block-diagonal,

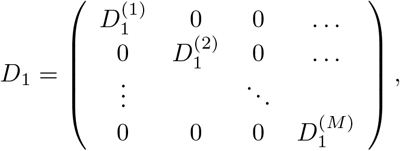

where 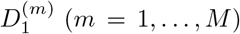 are square blocks along the diagonal of *D*_1_, and *M* is the total number of blocks. Such correlation structures are often approximately satisfied in genetic studies: for example, subjects are highly genetically correlated within the same family, and are remotely correlated across families. Let *s*_*m*_ denote the number of rows/columns in 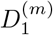, and *Y* ^(*m*)^ denote the *s*_*m*_ × 1 subvector of *Y* corresponding to the *m*^*th*^ block. For example, *Y* ^(1)^ is the subvector corresponding to the first *s*_1_ elements of *Y*. By construction, *Y* ^(*m*)^’s are independent and normally distributed with zero mean and covariance 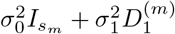, for *m* = 1,…,*M*.

We proceed with first establishing consistency and asymptotic normality of the HE estimator. By (3), the HE estimate is the ordinary least squares solution to a linear regression problem. For sparse block-diagonal *D*_1_, we simplify 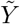 and 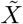 in (3) by discarding elements corresponding to zero entries of *D*_1_:

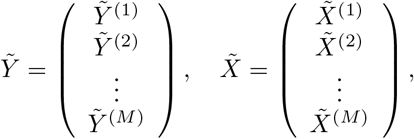

where we denote 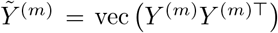 and 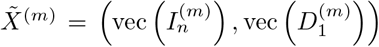, for *m* = 1, …, *M*. The independence of *Y* ^(*m*)^’s implies independence of 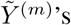. Thus, the HE estimator falls within the framework of generalized estimating equations with *I*_*n*_ as the *working correlation matrix*. The consistency and asymptotic normality properties of such an estimator are well known (see 32, Example 2.1 and 5.1). Specifically, as the number of blocks *M* →∞, regardless of whether the maximum block size 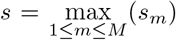 goes to infinity or not, it follows from Example 2.1 in [32] that the H<u>E</u> estimates 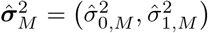 converge in probability to the true variance components 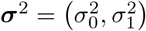 at a rate of 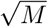 or faster. Moreover, when the maximum block size *s* does not go to infinity too fast as *M* →∞, the HE estimates are asymptotically normal, with a rate of 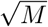 [32]:

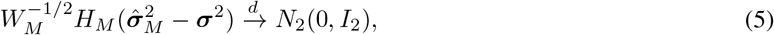

where

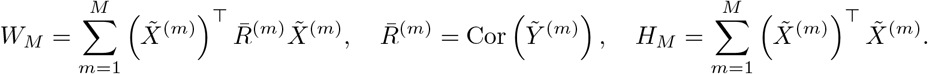

In genetic studies, it is typical that subjects belong to small unrelated groups so that s is bounded, and the number of groups *M* increases with increasing sample sizes. These settings satisfy the conditions for the HE estimator’s consistency and asymptotic normality.

We are now ready to establish the asymptotic properties of the REHE estimator, 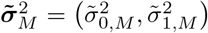. As implied by (4), the REHE estimates are different from the HE estimates 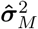 only when HE yields negative estimates for some variance components. Let 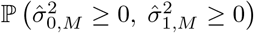 denote the probability of the HE estimates being non-negative, which equals 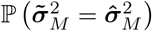. By (5), we have:

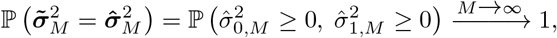

indicating asymptotic equivalence of the HE and REHE estimators. Note that despite their asymptotic equivalence, our simulation results in Section 4 clearly show the advantages of REHE over HE in finite samples. Next, we show that 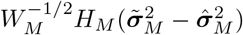 is *o_p_*(1):

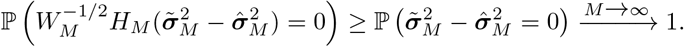

We then have

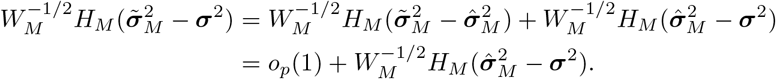

Therefore, we have established that the REHE estimator 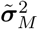 satisfies:

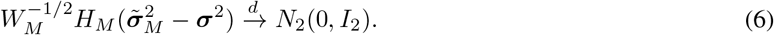

#### 2.2.2 REHE with resampling (reREHE)

Estimates obtained from REHE are non-negative. However, in some applications, a zero estimate for the variance component may still make the interpretation and/or subsequent analyses challenging. To address this issue, we equip REHE with a resampling procedure that provides strictly positive variance component estimates with high probability. The resampling procedure utilizes repeated subsamples, which can further improve the computational efficiency of REHE. The resulting approach is termed reREHE.

##### Algorithm 1 reREHE Approach Estimation.

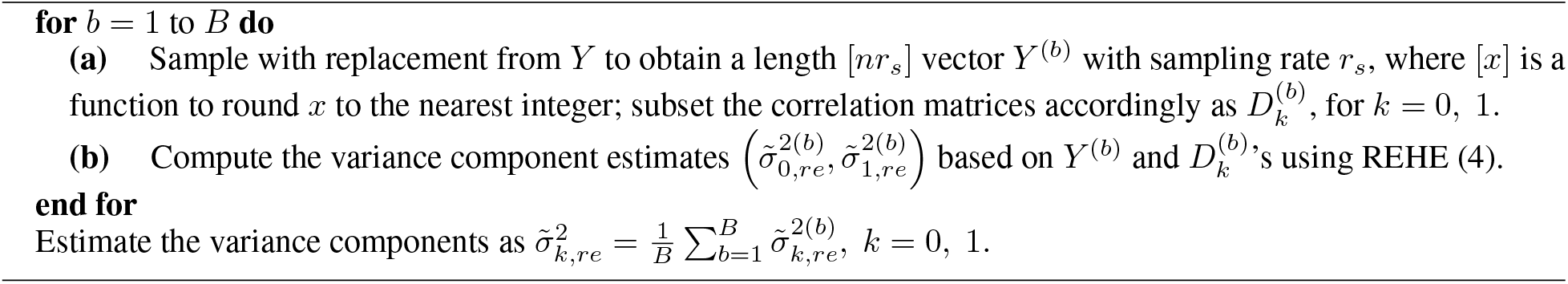

The idea of subsampling for REHE is simple: instead of estimating the variance components based on all the observations at the same time, we only use a small subsample of the data. The full-sample-based estimates and inference are usually well approximated by statistics based on subsamples [24]. Similar subsampling techniques are also extensively used in stochastic gradient descent methods [22]. Recently, subsampling has also been used with HE-based estimating equations [33]. Our proposed reREHE procedure described in Algorithm 1 is unique in that it subsamples repeatedly to obtain the estimates. Although at a cost of reduced computational efficiency compared to using a single subsample, this resampling offers considerable advantages. By averaging the estimates from repeated subsamples, the reREHE estimates have much higher accuracy compared to estimates based on a single subsample. At the same time, we obtain strictly positive estimates, unless in extremely rare cases when all subsamples yield zero estimates. Other summaries, such as median, can also be used to summarize estimates from repeated subsamples. When the sampling rate *r*_*s*_ and the number of subsamples *B* satisfy 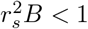, reREHE achieves higher computational efficiency than REHE. On the other hand, choosing larger *r*_*s*_ and *B* results in more stable results. In the simulation studies and data applications, we chose *B* = 50 and varied *r*_*s*_ within (0.05, 0.1) to achieve a balance between accuracy and computational efficiency.

#### 2.2.3 REHE with Fixed Effects

The REHE estimation procedure can be easily modified to accommodate fixed effects. Consider the full model (1) where *X* is the design matrix with *p* covariates and *β* is the fixed effect coefficient vector. Let 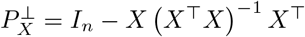 denote the projection matrix onto the orthogonal complement of the column space of *X*. We project the outcome *Y* and the random effects *γ*_*k*_’s (including the noise term *γ*_**0**_) as

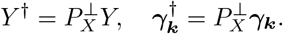

Recall that each random effect *γ*_*k*_ follows a normal distribution with zero mean and covariance *D*_*k*_. Writing 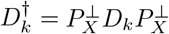, model (1) becomes

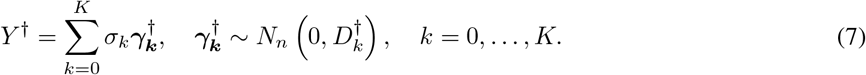

With model (7), we can directly apply the REHE approach as introduced in Section 2.2 to estimate the variance components. When the sample size *n* is large, computing the projected correlation matrices 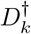 is time-consuming. In genetic and genomics applications, the number of fixed effect covariates *p* is much smaller than the sample size *n*. We also have balanced design in many of these applications. In those settings we are able to obtain good estimates of the fixed effects, and we can directly use the original correlation matrices *D*_*k*_ instead of projected matrices 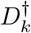. In such cases, as in our data applications and simulation studies, we expect results based on 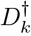 to be very close to those based on *D*_*k*_. We thus suggest using *D*_*k*_ for computational efficiency when estimating model (7) via REHE or reREHE.

If the fixed effect coefficients *β* are of interest, they can be estimated using ordinary least squares as 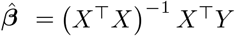, or weighted least squares as 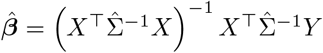, where 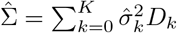 is based on previously estimated variance components 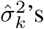. While the resulting 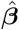 is consistent for *β*, one can iteratively update 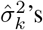 and 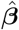 as in [29].

#### 2.2.4 Constructing Confidence Intervals with REHE

To obtain confidence intervals for variance component estimates, we use a parametric bootstrap procedure as summarized in Algorithm 2.

##### Algorithm 2 Parametric Bootstrap Confidence Interval Construction for REHE.

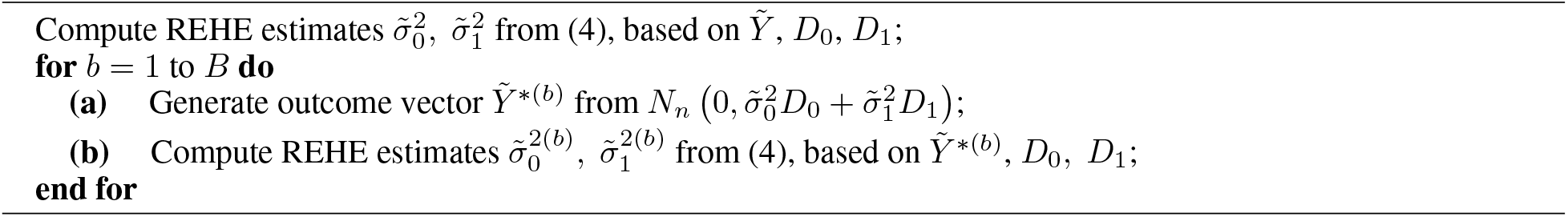

Using the bootstrap samples of REHE estimates, 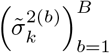, for *k* = 0, 1, we can construct Wald-type confidence intervals as

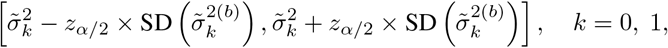

where *z*_*α/*2_ is the (1 – *α/*2) × 100th percentile of the standard normal distribution, and SD(·) denotes the sample standard deviation over bootstrap samples. Wald-type confidence intervals are valid, provided that the estimates are normally distributed. We can also construct quantile bootstrap confidence intervals as

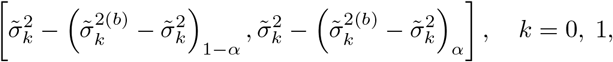

where 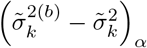 is the (1 − *α*) × 100th empirical quantile of 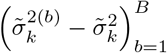.

For small sample sizes, the quantile confidence intervals are expected to be more robust than their Wald-type counterparts. When the REHE estimator is close to be normally distributed under large sample size, Wald-type confidence interval might have higher accuracy based on the same number of bootstrap samples. In simulation studies and real data applications, we chose the number of bootstrap samples *B* = 50 to balance between computational time and confidence interval accuracy. Confidence intervals for functions of variance components, such as heritability, can be similarly obtained by transforming the bootstrap samples accordingly.

When the correlation matrix *D*_1_ is dense and the sample size *n* is large, it is computationally prohibitive to compute a matrix decomposition (through Cholesky or singular value decomposition) of *D*_1_, which is required for the sampling step (a) in the Algorithm 2. To speed up the inference procedure in this case, we use correlation matrix sparsification [9]. The sparsification approximates the dense correlation matrix *D*_1_ by a sparse block-diagonal matrix 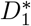, and thus largely accelerates matrix decomposition. The application of matrix sparsification to variance component inference in genetic studies is novel and is discussed in detail in Supplementary Note 1.2. Related simulation studies are shown in Supplementary Note 2.1 and Note 2.3.

## 3 Applications

### 3.1 GWAS and Heritability Study with HCHS/SOL Data

To evaluate the performance of REHE and reREHE in genetics applications, we conducted a genome-wide association analysis as well as a heritability analysis using a publicly available data set from the Hispanic Community Health Study/Study of Latinos (HCHS/SOL) [30, 4]. Before preprocessing, the selected HCHS/SOL data set contained 12,803 subjects with 4,100,028 single nucleotide polymorphisms (SNPs).

In the genome-wide association analysis, we tested the association between each SNP and the red blood cell count using a linear mixed model [4]. We first fit a null model, which was a linear mixed model without any genotype effect [1]. We included fixed effects covariates age, gender, cigarette use, field center indicator, genetic subgroup indicator, the first five principle components for population stratification effect, and individual sampling weights [4]. We removed subjects that have missing values for the above covariates, and included 12,502 subjects in the analysis. Relatedness among subjects was modelled by three random effects: genetic relatedness represented by kinship, membership of household, and membership of community group [4].

We separately applied REHE, reREHE and REML to estimate the null model. For reREHE, we chose sampling rate *r*_*s*_ = 0.1 and used mean summary function based on 50 repeated subsamples. With *n* =12,502 subjects to be analyzed, REHE only took 2.4 minutes to estimate the null model, a speed up of more than 10 folds compared to REML. reREHE was similarly fast as REHE. Based on each estimated null model, we applied score tests for the association of each SNP and the red blood cell count [4], and compared the resulting *p*-values. We focus here on genes with *p*-values no larger than 5 × 10^−8^ by at least one approach [5]. As shown in Figure 1a and 1b, results based on REHE and reREHE have negligible differences from those based on REML. This concordance among REML, reREHE and REHE is not surprising as the estimated variance components are similar (Figure 1c).

**Figure 1:**
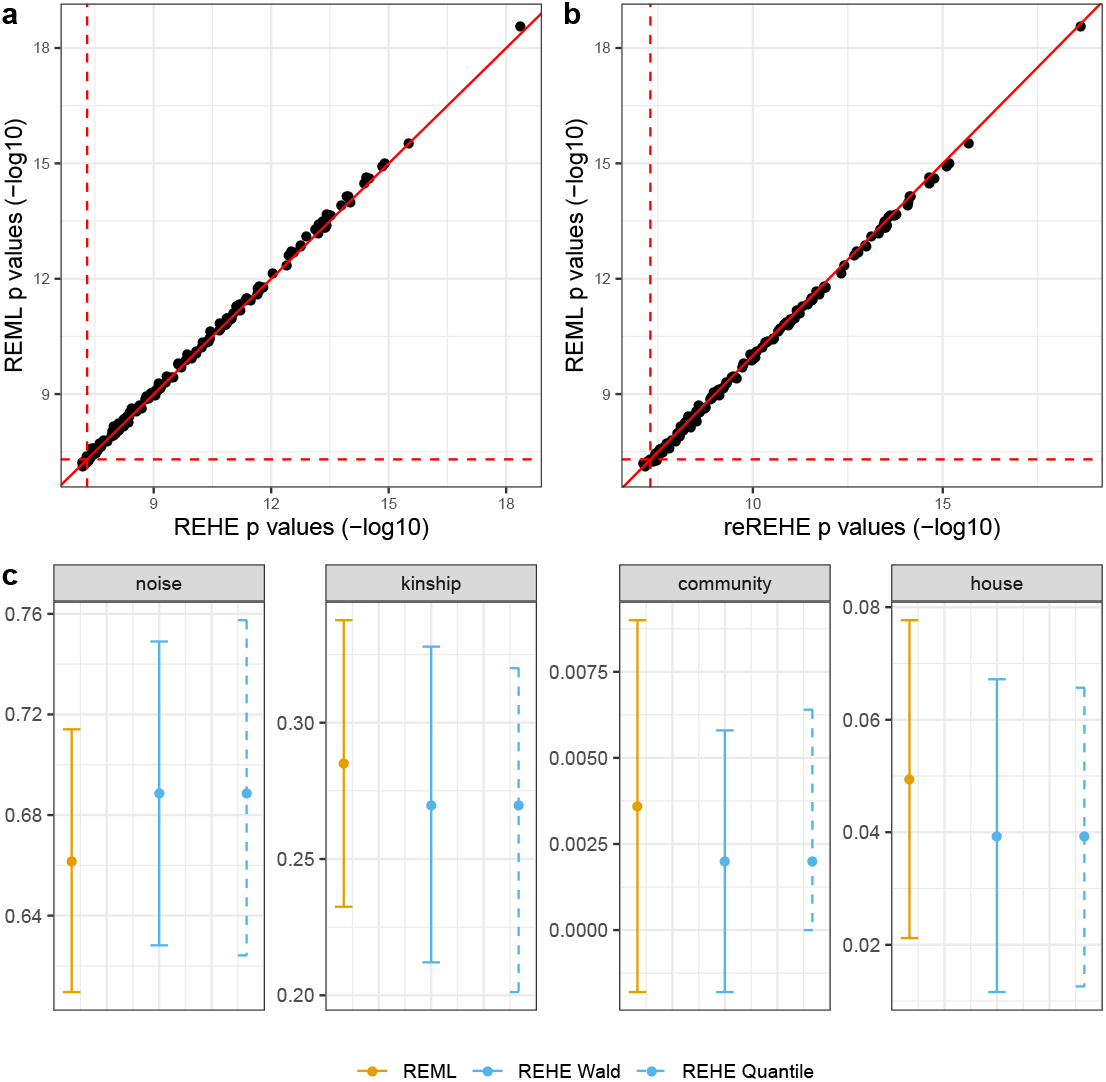
Results of genome-wide association testing analysis and heritability analysis with a HCHS/SOL data set. Gene association score test *p*-values (− log_10_ scale) based on: **a** - REML against REHE estimated null models; **b** - REML against reREHE estimated null models. Only genes with resulted *p*-values no larger than 5 × 10^−8^ by at least one approach were presented. **c** - Dots represent point estimates of proportion of variance attributed to noise, kinship (heritability), community membership and household membership; bars represent corresponding confidence intervals. Results by REHE and REML are displayed. Two types of REHE-based confidence intervals are presented: Wald-type confidence intervals (REHE Wald), and quantile-type confidence intervals (REHE Quantile).

For the heritability analysis, we used the same data set and fitted the same linear mixed null model as in the above genome-wide association analysis. The model was estimated based on REML and REHE separately. We obtained point estimates and confidence intervals for heritability (corresponding to the kinship correlation matrix), and proportions of variance explained by household membership, community block membership, and noise. REHE took 18.2 minutes to conduct the inference, compared to 23.9 minutes by REML. Heritability and variance proportions estimates, as well as the confidence intervals obtained by the REHE approach are all very similar compared to those obtained by REML (Figure 1c).

We used the R package *GENESIS* (*v2*.*14*.*3*, 3) for REML and conducting genome-wide association analysis. All analyses were conducted on a computer with 2 × 6-core Intel Xeon CPU E5-2620 @ 2.00GHz 128GB RAM.

### 3.2 Network-based Pathway Enrichment Analysis with Breast Cancer Data

To further demonstrate that REHE and reREHE facilitate downstream analysis with fast variance component estimation, we performed a network-based pathway enrichment analysis, with a breast cancer data set from The Cancer Genome Atlas (TCGA) [31], preprocessed by [20]. The data set contains RNA-seq measurements for 2,598 genes from 100 genetic pathways, with 403 subjects from the ER positive subtype and 117 from the ER negative subtype.

Network-based pathway enrichment analysis tests for differential gene pathways associated with particular phenotypes under different conditions [19]. It assumes a linear mixed model for the relationship between gene expressions and the phenotype (see Supplementary Note 1.3 and 19 for details). Here, we compared the activities of 100 genetic pathways between the two ER groups. The ER group-specific gene networks — more specifically, the adjacency and influence matrices supplied to the linear mixed model — were estimated according to [19]. We estimated the variance components using REML, REHE and reREHE. For reREHE, we chose the sampling rate *r*_*s*_ = 0.1 for sampling the subjects, and additionally sampled gene entries within each subject with sampling rate 0.5 (see Supplementary Note 1.3 for details). The reREHE estimate was based on the mean of 50 repeated subsamples. After obtaining the variance components estimates, we tested for differences in the activity of each of the 100 genetic pathways [28].

We observed substantial improvement in computational efficiency of reREHE and REHE compared to REML: reREHE- and REHE-based analyses both took less than 2 minutes, whereas analysis with REML took over 1 hour. Comparing the resulting *p*-values, REHE and reREHE produce slightly more conservative *p*-values than REML (Figure 2). Moreover, REHE yields a zero estimate for the noise variance component, the reREHE estimate is 0.0120, while the REML estimate is 0.266. The corresponding network variance estimates are also quite different: 0.273 by REML, 0.534 by reREHE and 0.610 by REHE. This may be an evidence that the variation explained by the network is much larger than the variation from noise in the true model. As illustrated in our additional simulation studies in Supplementary Note 2.4, REML may yield unreliable estimates under similar settings. We should thus take extra caution when interpreting REML-based estimates and test results in this application.

**Figure 2:**
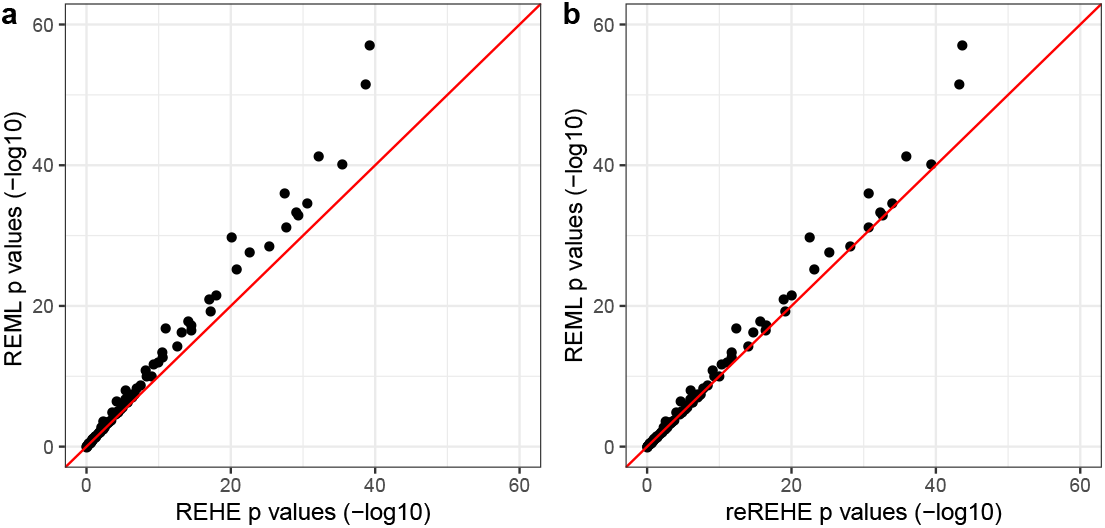
Results of network-based pathway enrichment analysis based on a breast cancer data set. *p*-values (− log_10_ scale) of t-tests for group difference of each gene pathway: **a** - compare REHE results against REML results; **b** - compare reREHE results against REML results. Two out-of-range data points are omitted from the plots, which correspond to: Glycosphingolipid biosynthesis - lacto and neolacto series pathway, with *p*-value 1.09 × 10^−307^ by REML, 5.67 × 10^−276^ by REHE, 3.74 × 10^−304^ by reREHE; and Caffeine metabolism pathway with *p*-value 3.97 × 10^−201^ by REML, 2.60 × 10^−179^ by REHE, and 2.13^−198^ by reREHE

We conducted all analyses using the R package *netgsa* (*v3*.*1*.*0*, 19) on a computer with 2 × 6-core Intel Xeon X5650 @ 2.67 GHz, 96GB RAM.

## 4 Simulation Studies

### 4.1 Simulation Settings

To benchmark the improvement of REHE and reREHE over HE and REML for variance components and heritability estimation, we generated synthetic data based on the HCHS/SOL design [30, 4].

reREHE and HE approaches were implemented only for point estimation comparison. We truncated negative HE estimates at zero. For reREHE, we used *B* = 50 repeated subsamples, and chose sampling rates *r*_*s*_ = 0.05 (reREHE 0.05) and *r*_*s*_ = 0.1 (reREHE 0.1). Point estimates were evaluated in terms of the root mean squared error (RMSE). We constructed Wald-type (REHE-Wald) and quantile-type (REHE-quantile) confidence intervals at 95% level for REHE estimates, and compared their performances with REML based confidence intervals in terms of coverage and interval width.

We simulated data based on the linear mixed model (2). We used sample size *n ∈* {3,000, 6,000, 9,000, 12,000}. For each sample size, we set the true values of the variance components to be 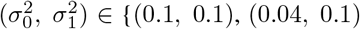, (0:1; 0:04), (0:01; 0:1), (0:1; 0:01)}. These values were chosen based on previous simulation study settings [29] and estimates from real data applications [6]. For each sample size *n* selected, we generate the correlation matrix *D*_1_ as a random sub-matrix of the kinship correlation matrix from the HCHS/SOL data set. For example, for *n* =3,000, we subsampled 3,000 out of 12,803 subjects without replacement, and used the corresponding (subsample) kinship correlation matrix as *D*_1_. Under each scenario, we ran 200 replicates. Computation was performed on a computer with 2 × 6-core Intel Xeon CPU E5-2620 @ 2.00GHz 128GB RAM.

We also conducted additional simulation studies to compare the approaches under different correlation structures; details of these experiments can be found in Supplementary Note 2.3.

### 4.2 Simulation Results

Simulation results clearly demonstrate the improvement in computational efficiency by REHE compared to REML. For point estimation, REHE was over 50 times faster than REML (Figure 3a). At the same time, REHE does not compromise estimation accuracy. Figure 3b and 3c show that REHE estimates of both the variance components and heritability are very close to those obtained by REML. Another advantage of REHE is that it corrects negative variance estimates from HE. To quantify this difference, We calculated the proportion of simulation replicates resulting in negative HE estimates (before zero-thresholding). This proportion reaches 23% with 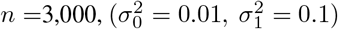, but reduces to 1.5% at *n* =12,000. As pointed out before, REHE automatically corrects the issue of negative estimates without hard-thresholding. Besides, REHE has lower RMSE for point estimates when HE estimates are likely being negative (Figure 3c).

**Figure 3:**
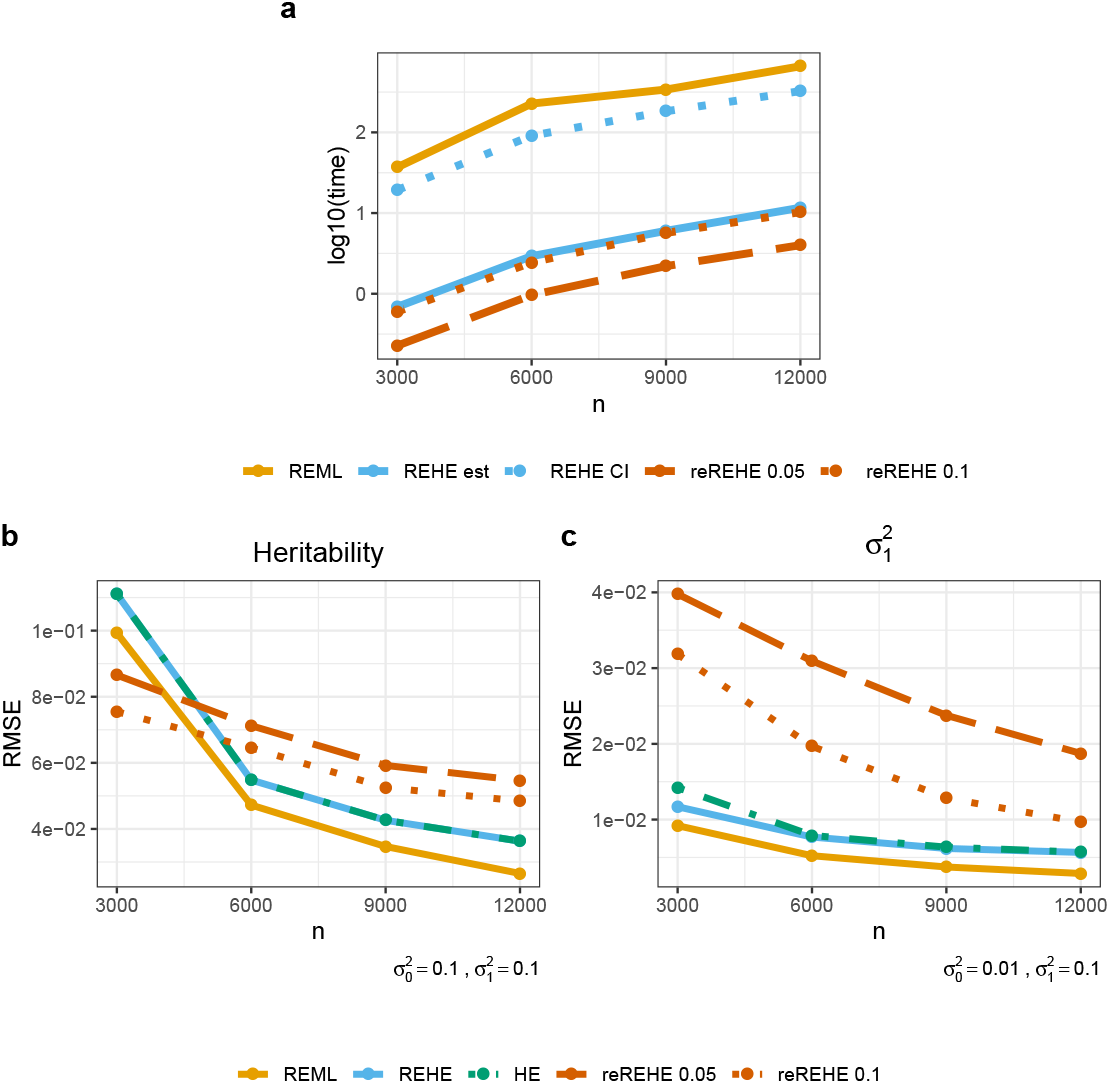
Simulation results for REHE and reREHE compared to REML. **a** - CPU time in seconds (log_10_ scale): time is presented separately for fitting the model using REML (REML), for only computing point estimates by REHE (REHE est), for constructing confidence interval for variance components with REHE (REHE CI), for only computing point estimates by reREHE with subsampling rate *r*_*s*_ = 0.05 (reREHE 0.05) and reREHE with subsampling rate *r*_*s*_ = 0.1 (reREHE 0.1). **b** - Root mean squared error (RMSE) for heritability estimation; simulation was based on true values 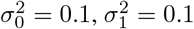, where 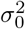 is the variance for noise, and 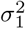 is the variance for the random effect. **c** - RMSE for 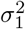 estimation; simulation was based on true values 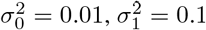.

The simulation results confirm that reREHE can provide strictly positive estimates with high probability. By providing a positive variance estimate where REHE gives a zero estimate (up to 23% of the simulation repetitions), reREHE is helpful for interpretation and downstream analysis, especially under small sample sizes. As shown in Figure 3b, reREHE based estimates have smaller RMSE than all other methods at sample size *n* =3,000. With larger samples, the RMSE of reREHE is comparable to other methods under some settings (Figure 3b), but is much larger in other settings (Figure 3c, Supplementary Note 2.1 and Note 2.3). Setting a higher subsampling rate (0.1 compared to 0.05) reduces RMSE (Figure 3b and 3c), but comes at the cost of reduced computational efficiency — reduction in computation time compared to REHE diminishes from 67% to 10% (Figure 3a).

Turning to inference of variance components and heritability, both REHE based quantile-type and Wald-type confidence intervals provide reasonably good coverage with comparable interval width to REML confidence intervals (Figure 4).

**Figure 4:**
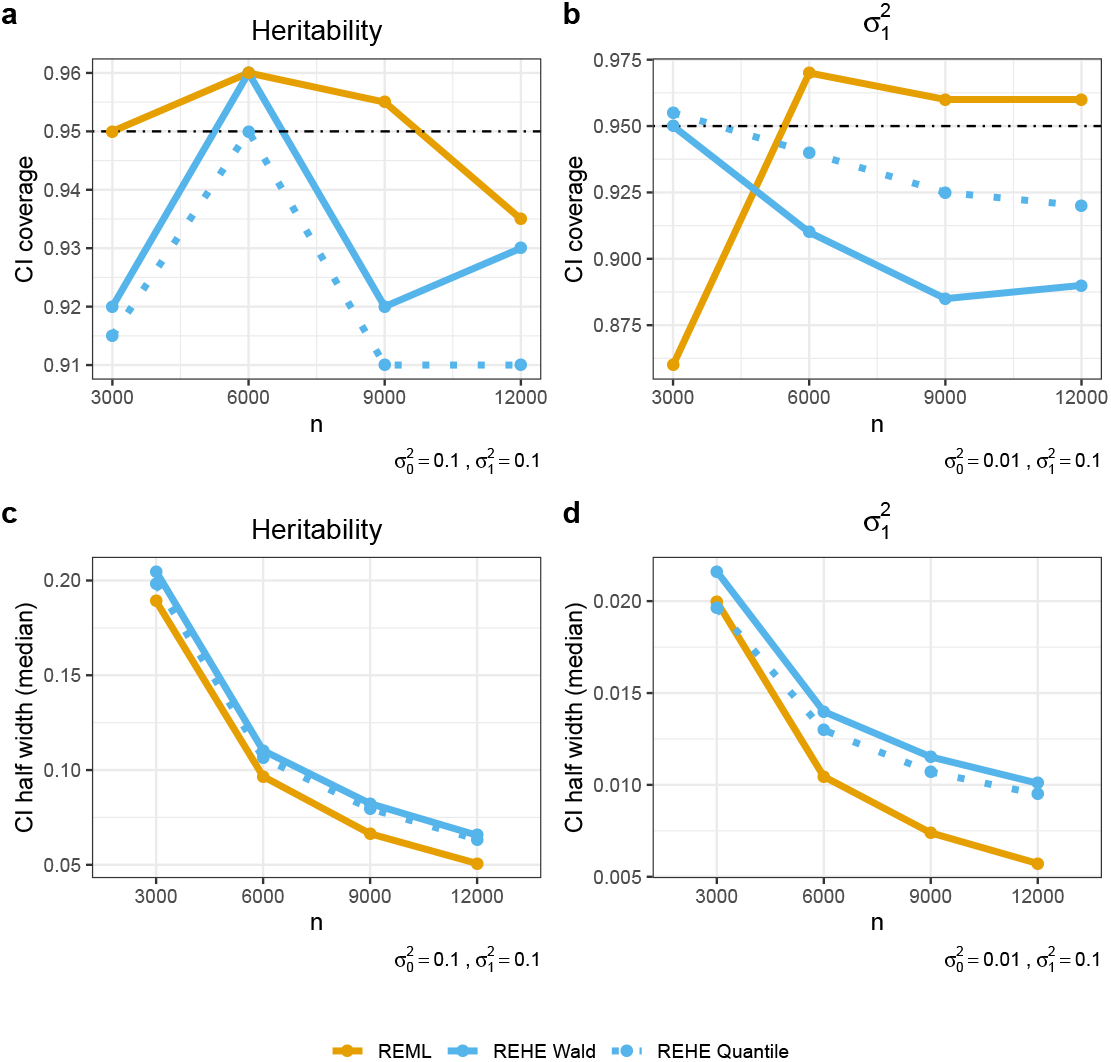
Simulation results for confidence interval performance in terms of coverage and width. 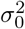 is the variance for noise, and 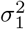 is the variance for the random effect. **a** - Coverage for heritability confidence interval (CI); simulation was based on true values 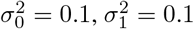. Monte Carlo error of 0.03 is expected for 200 simulation replications. **b** -Coverage for 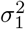 CI; simulation was based on true values 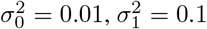. Monte Carlo error of 0.03 is expected for 200 simulation replicates. **c** - Line charts of median half width of heritability CI with increasing sample sizes; simulation was based on true values 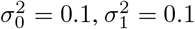. **d** - Line charts of median half width of 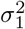 CI with increasing sample sizes; simulation was based on true values 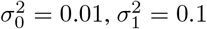.

The empirical coverage is close to nominal level under most cases (Figure 4a and 4b), considering a Monte Carlo error of 0.03 based on 200 simulation replicates. REHE quantile-type intervals generally have better coverage than Wald-type intervals when the true variance components are very different (Figure 4b). In terms of confidence interval width, quantile-type REHE confidence intervals are generally narrower than Wald-type, and both are comparable to REML-based intervals (Figure 4c and 4d). Inference based on REHE is more time-consuming than REHE-based point estimation; however, it still achieves 50% reduction of computation time compared to REML (Figure 3a).

Finally, in some simulation settings, we noticed that REML confidence intervals may suffer from under-coverage. For instance, with *n* =3,000 samples, when one variance component is substantially smaller, the coverage of REML confidence intervals drop below 87% (Figure 4b). In other settings, REML confidence intervals even have coverage below 60%, and have little improvement with increasing sample size (Supplementary Note 2.3). Another concern is the numerical stability of REML: REML fails to provide a confidence interval if the estimate of any variance component becomes zero during the iterative updates. We noticed frequent occurrence of this issue when the true variance components are unbalanced and the sample size is small. For *n* =3,000 and 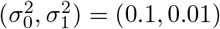, REML is unable to provide a confidence interval in 17.5% of the replicates. This proportion increases to 30.5% in other settings (Supplementary Note 2.3). We view these two issues as a warning sign for REML-based inference in real applications, especially when the underlying variance components are very different and the sample size is small. In contrast, REHE-based inference is robust across different settings with valid confidence intervals and acceptable empirical coverage.

The above simulation results are supported by evidence from additional simulation studies in Supplementary Note 2.3.

## 5 Discussion

We proposed REHE for fast estimation of variance components in linear mixed models. Through simulation studies and data applications, we demonstrated its substantial gain in computational efficiency over REML, with little compromise in point estimation accuracy. Compared to HE, REHE corrects the issue of negative estimates, and potentially has large gains in estimation accuracy. Therefore, REHE can be superior compared to HE and a good alternative to REML for point estimation of variance components in linear mixed models.

We also proposed reREHE based on the resampling technique to produce strictly positive variance component estimates with high probability in practice. Strictly positive estimates are more interpretable and may be more appealing for downstream analyses. Though reREHE estimates may have lower accuracy than REHE, the magnitude of the increase in RMSE is small in our experiments. We have also seen in the real data application that based on a subsampling rate of 0.1 and 50 subsamples, reREHE-based downstream analysis results are close to REML-based results. With suitably chosen subsampling rate and number of subsampling replicates, reREHE can achieve higher computational efficiency than REHE.

As mentioned previously, one can also use the median of the subsample results as the reREHE estimate. We explored this choice in the Supplementary Note 2.2 and Note 2.3. When the underlying variance components are very different, median-based reREHE estimates generally have smaller RMSE; otherwise mean-based reREHE performs better. However, median-based reREHE is more likely to yield zero estimates. A post hoc selection of the summary function can be made after observing the distribution of the subsample estimates based on reREHE.

As illustrated in the genome-wide association and pathway enrichment analysis examples, in many applications, only variance component estimates are needed for downstream analyses. The computation burden of REML-based estimation prohibits these analyses on large data sets. Restricting analysis to subsets of data reduces reliability and may yield contradictory conclusions. Given the fast and reliable estimates by REHE and reREHE in large data sets, we see great potential for their application in areas that only require point estimation of variance components.

When confidence intervals are also of interest, REHE remains a competitive alternative to REML for its robustness and numerical stability. As illustrated in our simulation studies, when the sample size is small and the true variance components are unbalanced, REML based inference is likely to suffer from numerical instability and/or poor coverage. REHE consistently provides valid inference across all settings. Therefore, when the true variance components are expected to be very different and the sample size is not large, we recommend REHE over REML if inference on variance components is needed, as REML-based inference results may be unreliable.

Constructing confidence intervals for variance components when sample size is large (e.g. *n >*10,000) is inherently computationally challenging. REHE only offers marginal improvements in computational efficiency over REML when it comes to inference. However, we can improve the computational efficiency for REHE confidence interval by parallelizing the bootstrap procedure. An alternative acceleration approach is to use correlation matrix sparsification (9, Supplementary Note 1.2). In the Supplementary Note 2.1 and Note 2.3, we explored the application of sparsification for constructing confidence intervals. Our conclusion is that sparsification improves computational efficiency in large sample settings (*n >*12,000); however, it may result in less robust confidence intervals for both REHE and REML. We did not explore application of sparsification to linear mixed models with more than one random effect. We expect a much larger sample size beyond which sparsification would show improvement in computational efficiency.

We did not explore confidence interval construction for reREHE estimates in this paper. Due to the repeated subsampling procedure of reREHE, an analytical expression for the confidence intervals is not trivial. The parametric bootstrap procedure for REHE confidence interval construction is readily extendable to reREHE, which we expect to have similar performance as REHE confidence intervals. However, the computation burden will also be similar to those of REHE confidence intervals. Future research should explore fast inference procedure for REHE and reREHE estimates.

## 6 Software

The proposed methods are implemented with codes written in the R language, which are available at https://github.com/yuek9/REHE.

## Supplement

Supplementary material contains additional information on REHE, reREHE and simulation studies. It is available online at https://www.biorxiv.org/.

## References

[1] Yurii S Aulchenko, Dirk Jan De Koning, and Chris Haley. Genomewide rapid association using mixed model and regression: a fast and simple method for genomewide pedigree-based quantitative trait loci association analysis. Genetics, 177(1):577–585, 2007.

[2] Andreas Weingessel Berwin A. Turlach. quadprog: Functions to Solve Quadratic Programming Problems, 2019. R package version 1.5-7.

[3] Matthew P. Conomos, Stephanie M. Gogarten, Lisa Brown, Han Chen, Ken Rice, Tamar Sofer, Timothy Thornton, and Chaoyu Yu. GENESIS: GENetic EStimation and Inference in Structured samples (GENESIS): Statistical methods for analyzing genetic data from samples with population structure and/or relatedness, 2019. R package version 2.14.3.

[4] Matthew P Conomos, Cecelia A Laurie, Adrienne M Stilp, Stephanie M Gogarten, Caitlin P McHugh, Sarah C Nelson, Tamar Sofer, Lindsay Fernández-Rhodes, Anne E Justice, Mariaelisa Graff, et al. Genetic diversity and association studies in US Hispanic/Latino populations: applications in the Hispanic Community Health Study/Study of Latinos. The American Journal of Human Genetics, 98(1):165–184, 2016.

[5] Frank Dudbridge and Arief Gusnanto. Estimation of significance thresholds for genome wide association scans. Genetic Epidemiology: The Official Publication of the International Genetic Epidemiology Society, 32(3):227–234, 2008.

[6] Mogens Fenger. Heritability and genetics of lipid metabolism. Future Lipidology, 2(4):433–444, 2007.

[7] Vojtečh Franc, Václav Hlaváč, and Mirko Navara. Sequential coordinate-wise algorithm for the non-negative least squares problem. In International Conference on Computer Analysis of Images and Patterns, pages 407–414. Springer, 2005.

[8] Arthur R Gilmour, Robin Thompson, and Brian R Cullis. Average information REML: an efficient algorithm for variance parameter estimation in linear mixed models. Biometrics, 55:1440–1450, 1995.

[9] Stephanie M Gogarten, Tamar Sofer, Han Chen, Chaoyu Yu, Jennifer A Brody, Timothy A Thornton, Kenneth M Rice, and Matthew P Conomos. Genetic association testing using the genesis r/bioconductor package. Bioinformatics, 35(24):5346–5348, 2019.

[10] Donald Goldfarb and Ashok Idnani. A numerically stable dual method for solving strictly convex quadratic programs. Mathematical Programming, 27(1):1–33, 1983.

[11] Franklin A Graybill. On quadratic estimates of variance components. The Annals of Mathematical Statistics, 25(2):367–372, 1954.

[12] Franklin A Graybill and Robert A Hultquist. Theorems concerning eisenhart’s model ii. The Annals of Mathematical Statistics, 32(1):261–269, 1961.

[13] Franklin A Graybill and AW Wortham. A note on uniformly best unbiased estimators for variance components. Journal of the American Statistical Association, 51(274):266–268, 1956.

[14] JK Haseman and RC Elston. The investigation of linkage between a quantitative trait and a marker locus. Behavior Genetics, 2(1):3–19, 1972.

[15] Longda Jiang, Zhili Zheng, Ting Qi, Kathryn E Kemper, Naomi R Wray, Peter M Visscher, and Jian Yang. A resource-efficient tool for mixed model association analysis of large-scale data. Nature Genetics, 51(12):1749– 1755, 2019.

[16] Hyun Min Kang, Jae Hoon Sul, Susan K Service, Noah A Zaitlen, Sit-yee Kong, Nelson B Freimer, Chiara Sabatti, Eleazar Eskin, et al. Variance component model to account for sample structure in genome-wide association studies. Nature Genetics, 42(4):348–354, 2010.

[17] Dongmin Kim, Suvrit Sra, and Inderjit S Dhillon. A new projected quasi-newton approach for the nonnegative least squares problem. Technical Report TR-06-54, Computer Science Department, University of Texas at Austin Austin, 2006.

[18] Charles L Lawson and Richard J Hanson. Solving least squares problems, volume 15. Siam, 1995.

[19] Jing Ma, Ali Shojaie, and George Michailidis. Network-based pathway enrichment analysis with incomplete network information. Bioinformatics, 32(20):3165–3174, 2016.

[20] Jing Ma, Ali Shojaie, and George Michailidis. A comparative study of topology-based pathway enrichment analysis methods. BMC bioinformatics, 20(1):546, 2019.

[21] Kaarina Matilainen, Esa A Mäntysaari, Martin H Lidauer, Ismo Strandén, and Robin Thompson. Employing a monte carlo algorithm in newton-type methods for restricted maximum likelihood estimation of genetic parameters. PloS one, 8(12):e80821–e80821, 2013.

[22] Michael R Metel. Mini-batch stochastic gradient descent with dynamic sample sizes. arXiv preprint arXiv:1708.00555, 2017.

[23] H Desmond Patterson and Robin Thompson. Recovery of inter-block information when block sizes are unequal. Biometrika, 58(3):545–554, 1971.

[24] Dimitris N Politis, Joseph P Romano, and Michael Wolf. Subsampling. Springer Science & Business Media, 1999.

[25] C Radhakrishna Rao. Estimation of heteroscedastic variances in linear models. Journal of the American Statistical Association, 65(329):161–172, 1970.

[26] Dieter Rasch and O Masata. Methods of variance component estimation. Czech Journal of Animal Science, 51(6):227, 2006.

[27] Shayle R Searle. An overview of variance component estimation. Metrika, 42(1):215–230, 1995.

[28] Ali Shojaie and George Michailidis. Analysis of gene sets based on the underlying regulatory network. Journal of Computational Biology, 16(3):407–426, 2009.

[29] Tamar Sofer. Confidence intervals for heritability via Haseman-Elston regression. Statistical Applications in Genetics and Molecular Biology, 16(4):259–273, 2017.

[30] Paul D Sorlie, Larissa M Avilés-Santa, Sylvia Wassertheil-Smoller, Robert C Kaplan, Martha L Daviglus, Aida L Giachello, Neil Schneiderman, Leopoldo Raij, Gregory Talavera, Matthew Allison, Lisa Lavange, Lloyd E Chambless, and Gerardo Heiss. Design and implementation of the hispanic community health study/study of latinos. Annals of Epidemiology, 20(8):629–41, Aug 2010.

[31] TCGA. Comprehensive molecular portraits of human breast tumours. Nature, 490(7418):61, 2012.

[32] Minge Xie and Yaning Yang. Asymptotics for generalized estimating equations with large cluster sizes. The Annals of Statistics, 31(1):310–347, 2003.

[33] Xiang Zhou. A unified framework for variance component estimation with summary statistics in genome-wide association studies. The Annals of Applied Statistics, 11(4):2027, 2017.

