## Supplementary Materials for "REHE: Fast Variance Components Estimation for Linear Mixed Models"

A PREPRINT

**Kun Yue**
Department of Biostatistics
University of Washington
Seattle, WA, USA

**Jing Ma**
Public Health Sciences Division
Fred Hutchinson Cancer Research Center
Seattle, WA, USA

**TIMOTHY THORNTON**
Department of Biostatistics
University of Washington
Seattle, WA, USA

**Ali Shojaie***
Department of Biostatistics
University of Washington
Seattle, WA, USA


February 3, 2021

### Supplementary Note 1: Extensions of REHE

#### 1.1 Analytical Solutions to REHE

With only two variance components in the linear mixed model ($K = 1$), there is a closed form solution to (2.4) in the main paper for REHE variance component estimates. Equation (2.4) in the main paper can be reformulated as:

$$(x, y) = \arg\min_{x,y} \; ax^2 + by^2 + cxy + dx + ey + f$$
$$\text{subject to } x, y \geq 0, \; a, b > 0, \; 4ab \neq c^2.$$

The condition $4ab \neq c^2$ corresponds to matrix $\tilde{X}^\top \tilde{X}$ being invertible, where $\tilde{X}$ is introduced in the main paper Section 2.1 as the design matrix in the HE regression. The closed form solution is:

$$(x, y) = \begin{cases} (\frac{ce - 2bd}{4ab - c^2}, \frac{cd - 2ae}{4ab - c^2}), & \text{if } \frac{ce - 2bd}{4ab - c^2} \geq 0, \; \frac{cd - 2ae}{4ab - c^2} \geq 0; \\ (0, -\frac{e}{2b}), & \text{if } \frac{ce - 2bd}{4ab - c^2} < 0, \; \frac{e}{2b} \leq 0; \\ (-\frac{d}{2a}, 0), & \text{if } \frac{cd - 2ae}{4ab - c^2} < 0, \; \frac{d}{2a} \leq 0; \\ (0, 0), & \text{if } \frac{d}{2a} > 0, \; \frac{e}{2b} > 0. \end{cases} \tag{S1}$$

#### 1.2 REHE Confidence Interval Construction with Sparse Correlation Matrix

When the sample size $n$ is large and the correlation matrix $D_1$ is dense, constructing REHE confidence intervals is computationally demanding. The main burden comes from matrix decomposition when sampling $\tilde{Y}^*$ from $N_n(0, \Sigma)$ in step (a) of Algorithm 2 in the main paper. Subsample $\tilde{Y}^*$ is commonly obtained by computing $\Sigma^{1/2}e$, where $\Sigma = \Sigma^{1/2}(\Sigma^{1/2})^\top$ and $e$ sampled from $N_n(0, I_n)$. Matrix $\Sigma^{1/2}$ is usually obtained by Cholesky decomposition or

*correspondence author

singular value decomposition, which requires $O(n^3)$ operations. We propose to accelerate this procedure through approximating the dense correlation matrix $D_1$ with a sparse block-diagonal matrix $D_1^*$ [2]. In genetics studies, it is reasonable to assume high relatedness within small groups of subjects and low relatedness across groups, such as for kinship relatedness and household membership. Such relatedness structure can be well-approximated by sparse block-diagonal correlation matrices, through thresholding the off-diagonal entries of the original dense correlation matrices and rearranging the rows/columns [2]. The resulting sparse block-diagonal correlation matrices can substantially speed up the matrix decomposition step, and thus the inference procedure. Bootstrap confidence intervals obtained with sparse $D_1^*$ are expected to have good coverage. We used the function *makeSparseMatrix* in the R package *GENESIS* (*v2.14.3*, 1) to implement the correlation matrix sparsification procedure. The sparsification threshold was set at $2^{-5.5}$, which corresponds to the fifth degree relatedness for the kinship matrix [2].

#### 1.3 REHE and reREHE for Network-based Pathway Enrichment Analysis

Network-based pathway enrichment analysis assumes the following linear mixed model for the association between gene expressions and the phenotype [3]:

$$Y_i^{(t)} = \Lambda_t \mu_t + \Lambda_t \gamma_i + \epsilon_i, \quad i = 1, \ldots, n_t, \quad t = 1, 2, \ldots T, \tag{S2}$$

where length $p$ vector $Y_i^{(t)}$ is the observed phenotype for the $i^{th}$ subject in group $t$, the group-specific network influence matrices $\Lambda_t$'s are treated as known from external sources, length $p$ vectors $\mu_t$'s denote the group-specific mean signal for the $p$ gene entries, random effects $\gamma_i$'s follow distribution $N(0, \sigma_1^2 I_p)$, error terms $\epsilon_i \sim N(0, \sigma_0^2 I_p)$, and $n_t$ denotes the sample size for group $t$, $t = 1, \ldots, T$. We can easily format this linear mixed model into the form of model (2.1) in the main paper. The resulting random effect correlation matrix $D_1$ has a block-diagonal structure:

$$D_1 = \begin{pmatrix} D_1^{(1)} & 0 & 0 & 0 & 0 & \ldots \\ 0 & D_2^{(1)} & 0 & 0 & 0 & \ldots \\ \vdots & & \ddots & & & \\ 0 & 0 & \ldots & D_{n_1}^{(1)} & 0 & \ldots \\ \vdots & & & & \ddots & \\ 0 & 0 & 0 & 0 & 0 & D_{n_T}^{(T)} \end{pmatrix},$$

where $D_i^{(t)} = \Lambda_t \Lambda_t^\top \ (i = 1, \ldots, n_t, \ t = 1, \ldots, T)$ are $p \times p$ square blocks corresponding to the $i^{th}$ subject in group $t$. Based on the special correlation structure of $D_1$, we can write out the expression for computing the REHE variance component estimates as:

$$(\tilde{\sigma}_0^2, \ \tilde{\sigma}_1^2) = \arg \min_{(\sigma_0^2 \geq 0, \ \sigma_1^2 \geq 0)} \left( \tilde{Y} - \tilde{X} \boldsymbol{\sigma^2} \right)^\top \left( \tilde{Y} - \tilde{X} \boldsymbol{\sigma^2} \right),$$

where

$$\tilde{Y} = \begin{pmatrix} \text{vec}\left(Y_1^{(1)} Y_1^{(1)\top}\right) \\ \text{vec}\left(Y_2^{(1)} Y_2^{(1)\top}\right) \\ \ldots \\ \text{vec}\left(Y_1^{(2)} Y_1^{(2)\top}\right) \\ \ldots \\ \text{vec}\left(Y_{n_T}^{(T)} Y_{n_T}^{(T)\top}\right) \end{pmatrix}, \quad \tilde{X} = \begin{pmatrix} \text{vec}\left(D_1^{(1)}\right) & \text{vec}\left(\mathbf{I}_p\right) \\ \text{vec}\left(D_2^{(1)}\right) & \text{vec}\left(\mathbf{I}_p\right) \\ \ldots & \\ \text{vec}\left(D_1^{(2)}\right) & \text{vec}\left(\mathbf{I}_p\right) \\ \ldots & \\ \text{vec}\left(D_{n_T}^{T}\right) & \text{vec}\left(\mathbf{I}_p\right) \end{pmatrix}, \quad \boldsymbol{\sigma^2} = \begin{pmatrix} \sigma_0^2 \\ \sigma_1^2 \end{pmatrix}.$$

When applying the reREHE approach to NetGSA, in addition to subsampling the $n$ observations, we can sample the $p$ gene entries. The procedure is described in Algorithm S1.

### Supplementary Note 2: Additional Simulation Results

#### 2.1 Simulation Results on Matrix Sparsification

Correlation matrix sparsification accelerates confidence interval construction with REHE (Supplementary Note 1.2), and can also benefit computation with REML [2]. We conducted a simulation study to apply sparsification to both

**Algorithm S1** reREHE Approach for NetGSA.

**for** $b = 1$ to $B$ **do**

**(a)** Sample $[nr_s]$ out of $n$ subjects with replacement with sampling rate $r_s$, where $[x]$ denotes rounding $x$ to the nearest integer. The corresponding observation vectors are $Y_{i'},\ i' = 1, \ldots, [nr_s]$;

**(b)** For each observation vector $Y_{i'}$ in the subsample, randomly sample $[pr_p]$ out of $p$ gene entries with replacement with sampling rate $r_p$. The resulted observation vector $Y^*_{i'}$ is of dimension $[pr_p]$ by 1, and the outcome vector $Y^{*(b)}$ for the whole subsample is of length $[nr_s][pr_p]$.

**(c)** Subset the correlation matrix $D_1$ accordingly as $D_1^{*(b)}$: first only keep blocks $D_1^{(i')}$ corresponding to the subsampled observations; then for each $D_1^{(i')}$, only keep the submatrix with rows and columns corresponding to the subsampled gene entries.

**(d)** Compute variance component estimations $(\tilde{\sigma}^{2(b)}_{0,re}, \tilde{\sigma}^{2(b)}_{1,re})$ with $Y^{*(b)}$ and $D_1^{*(b)}$ based on the REHE approach.

**end for**

Compute estimates of variance components as $\tilde{\sigma}^2_{k,re} = \frac{1}{B}\sum_{b=1}^{B} \tilde{\sigma}^{2(b)}_{k,re}, \quad k = 0,\ 1.$

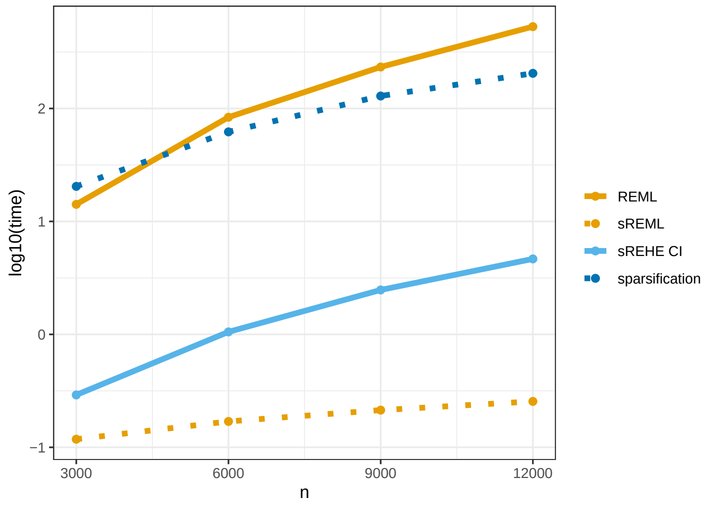


Figure S1: Results of the simulation study on matrix sparsification: Computation time (in seconds, $\log_{10}$ scale) for REML and REHE based on matrix sparsification, benchmarked with computation time for REML. We separately illustrate time for sparsifying the kinship matrix (sparsification), time for fitting the model with REML after obtaining the sparsified kinship matrix (sREML), and time for REHE confidence interval construction based on sparsified kinship matrix (sREHE CI).

REHE and REML for inference of variance components. We used the same HCHS/SOL design setting as had been used by the simulation study in the main paper Section 4. We compared their performances in terms of computation time, root mean squared error (RMSE) of variance component estimates, and confidence interval coverage and width. *sREML* refers to REML with the sparsified kinship correlation matrix. *sREHE Wald* and *sREHE Quantile* refers to, respectively, REHE Wald- and quantile-type confidence intervals based on sparsified kinship matrix. The REHE point estimation is still based on the original dense kinship matrix, because it is fast and more accurate than that obtained with sparsification.

Both REHE and REML inferences based on the sparse kinship matrix are substantially faster than those based on the original kinship matrix (Figure S1). We noticed that the sparsification procedure itself is computationally demanding. This may make sparsification less appealing when there are multiple dense correlation matrices to process, or when the sample size is not large enough. However, as the sample size grows, inference with REML becomes so slow that it can still be worthwhile to use matrix sparsification. At sample size $n$ =12,000, REHE inference with one sparsified correlation matrix (one random effect) reduces the REML computation with a dense correlation matrix by 50%.

The RMSE of point estimates by sREML is always larger than REML, and sometimes much larger than the RMSE of REHE estimates (Figure S2). The coverage of confidence intervals by methods based on sparsification is not as stable: sREHE shows conservative coverage, and sREML has coverage lower than 0.75 at sample size $n$ =3,000 (Figure S3). This illustrates that sparsification speeds up the computation but at the cost of inference accuracy. For the same method,

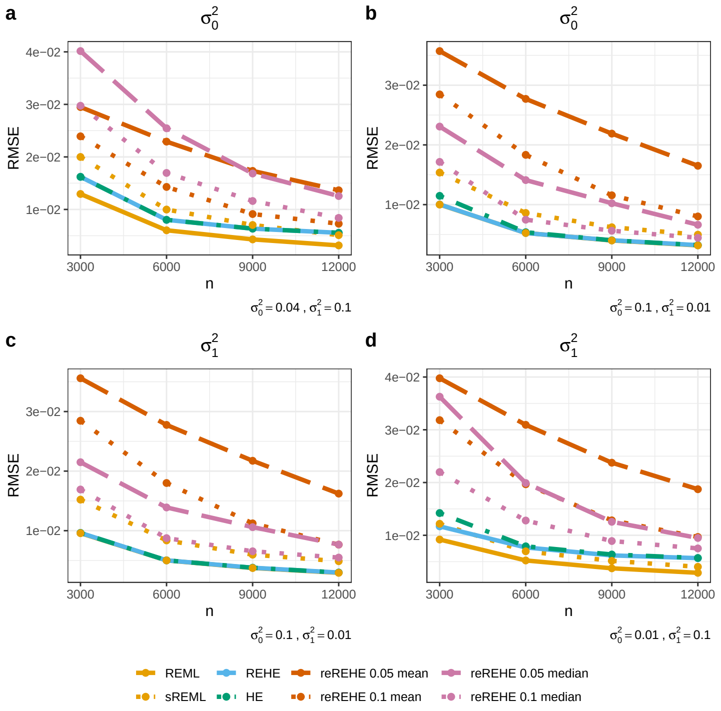


Figure S2: Results of the simulation study on matrix sparsification and on different reREHE summary functions: RMSE of point estimates by REML, REML with matrix sparsification (sREML), REHE, HE, and different implementations of reREHE: reREHE based on subsampling rate 0.05 and mean summary function (reREHE 0.05 mean), reREHE based on subsampling rate 0.1 and mean summary function (reREHE 0.1 mean), reREHE based on subsampling rate 0.05 and median summary function (reREHE 0.05 median), reREHE based on subsampling rate 0.1 and median summary function (reREHE 0.1 median). **a** - RMSE for $\sigma_0^2$ estimates, with true values $\sigma_0^2 = 0.04$, $\sigma_1^2 = 0.1$. **b** - RMSE for $\sigma_0^2$ estimates, with true values $\sigma_0^2 = 0.1$, $\sigma_1^2 = 0.01$. **c** - RMSE for $\sigma_1^2$ estimates, with true values $\sigma_0^2 = 0.1$, $\sigma_1^2 = 0.01$. **d** - RMSE for $\sigma_1^2$ estimates, with true values $\sigma_0^2 = 0.01$, $\sigma_1^2 = 0.1$.

confidence interval width is generally wider with sparsification than without (Figure S4). We found sREML and REHE yield comparable confidence interval width.

#### 2.2 Simulation Results on Different reREHE Summary Function

The reREHE approach can use different summary functions, such as mean and median functions, to summarize results of repeated subsamples. We implemented median based reREHE for comparison of estimation accuracy against mean based reREHE. We denote mean based reREHE results by *reREHE 0.05 mean* and *reREHE 0.1 mean*, and denote median based reREHE results by *reREHE 0.05 median* and *reREHE 0.1 median*, for subsampling rates 0.05 and 0.1 respectively. We used the same simulation setting as in Supplementary Note 2.1.

As illustrated by Figure S2, when the true variance components are very different (taking on values of 0.1 and 0.01), median based reREHE has smaller RMSE than mean based reREHE; otherwise mean based reREHE has smaller RMSE. Higher subsampling rate greatly improves estimation accuracy for both reREHE approaches. In general, reREHE approaches result in less accurate but acceptable estimates compared to REML and REHE results.

Mean based reREHE provides positive estimates with high probability, whereas median based reREHE is more likely to yield a zero estimate in practise. The trade-off among accuracy, interpretability and computational efficiency brought by reREHE will be left at user’s choice.

#### 2.3 Simulation Results on Heritability

We conducted additional simulation studies based on HCHS/SOL design, using three different artificial kinship matrices to compare REHE and reREHE against REML and HE. In the first two settings, we generated block-diagonal kinship

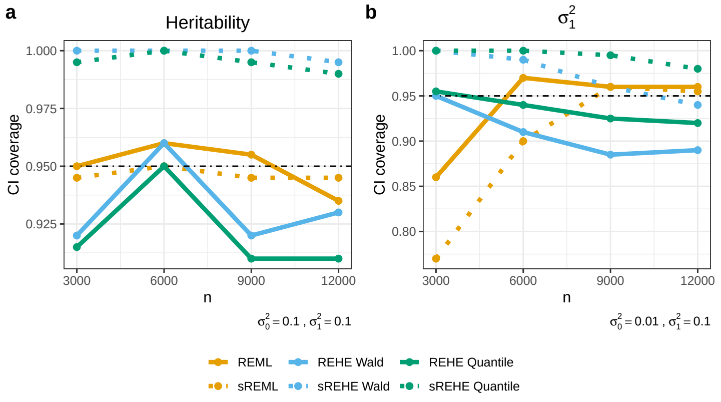


Figure S3: Results of the simulation study on matrix sparsification: confidence interval coverage with 200 replicates (Monte Carlo error approximately 0.03). We present results for confidence intervals constructed based on REML, REML with matrix sparsification (sREML), Wald-type intervals with REHE (REHE Wald), Wald-type intervals with REHE and matrix sparsification (sREHE Wald), quantile-type intervals with REHE (REHE Quantile), and quantile-type intervals with REHE and matrix sparsification (sREHE Quantile). **a** - Confidence interval coverage for heritability, with true values $\sigma_0^2 = 0.1$, $\sigma_1^2 = 0.1$. **b** - Confidence interval coverage for $\sigma_1^2$, with true values $\sigma_0^2 = 0.01$, $\sigma_1^2 = 0.1$.

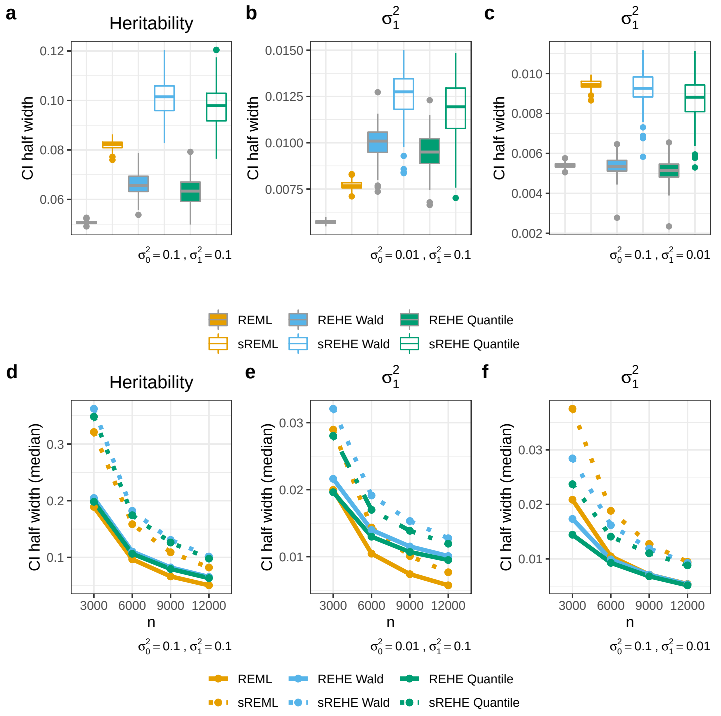


Figure S4: Results of the simulation study on matrix sparsification: boxplots of confidence interval half width, and line charts of median confidence interval half width against sample sizes. The boxplots follow standard Tukey representations, where center line represents median, box limits represent lower and upper quartiles, whiskers represent 1.5x interquartile range, and points represent outliers. We present results for confidence intervals constructed based on REML, REML with matrix sparsification (sREML), Wald-type intervals with REHE (REHE Wald), Wald-type intervals with REHE and matrix sparsification (sREHE Wald), quantile-type intervals with REHE (REHE Quantile), and quantile-type intervals with REHE and matrix sparsification (sREHE Quantile). The targeted parameter of the confidence interval is indicated by the title of each sub-plot. True values of variance components under each scenario ware presented on the bottom-right corner of each sub-plot.

matrix $D_1$ with $3 \times 3$ blocks $D_b$, such that $D_1$ had the form

$$D_1 = \begin{pmatrix} D_b & 0 & 0 & \dots \\ 0 & D_b & 0 & \dots \\ 0 & 0 & D_b & \\ \vdots & & & \ddots \end{pmatrix}.$$

Setting 1 represents a population with high correlation within groups, whereas setting 2 represents a population where subjects are remotely related even within groups. For setting 3, we generated a dense kinship matrix which is approximately block-diagonal, and then generated random values from $\mathrm{Unif}[-0.001, 0.001]$ to fill the off-diagonal zero entries in $D_1$. Setting 3 mimics the dense kinship matrix in real data applications, which may render estimation and inference more difficult. We set the values of $D_b$ in each setting as:

Setting 1: $$D_b = \begin{pmatrix} 1 & 0.8 & 0.2 \\ 0.8 & 1 & 0.4 \\ 0.2 & 0.4 & 1 \end{pmatrix};$$

Setting 2: $$D_b = \begin{pmatrix} 1 & 0.05 & 0.05 \\ 0.05 & 1 & 0.1 \\ 0.05 & 0.1 & 1 \end{pmatrix};$$

Setting 3: $$D_b = \begin{pmatrix} 1 & 0.1 & 0.05 \\ 0.1 & 1 & 0.3 \\ 0.05 & 0.3 & 1 \end{pmatrix}.$$

Based on each of the three kinship matrices, we ran 200 replicates for each combination of $(\sigma_0^2,\ \sigma_1^2) \in \{(0.1,\ 0.1), (0.04,\ 0.1), (0.1, 0.04), (0.01,\ 0.1), (0.1,\ 0.01)\}$ and $n \in \{3,000,\ 6,000,\ 9,000,\ 12,000\}$. We sparsified the kinship matrices when constructing confidence intervals for REHE and REML.

Computation time of each method is very similar to those presented in the main paper and in Supplementary Note 2.1 (Figure S1). Although the kinship matrix is already sparse and block-diagonal under setting 1 and 2, the sparsification algorithm is not able to process the matrices quickly. Sparsification procedures in these settings take similar amount of computation time as for sparsifying dense kinship matrices of the same size. This indicates room of improvement for the sparsification algorithm. We recommend the users to skip the sparsification procedure if the available correlation matrices are approximately sparse and block-diagonal such that computing their inverses are computationally efficient.

Figure S5 illustrates the RMSE. reREHE has similar performance as those described in Supplementary Note 2.1. The RMSE of sREML is always indistinguishable from that of REML. REHE estimates are close to REML estimates, and their RMSE often has negligible difference (e.g. Figure S5-b, S5-e, S5-h). When the true variance component values are different, improvement of REHE over HE is more pronounced than those illustrated in the main paper and Supplementary Note 2.1 (Figure S5-e, S5-f). Such improvement in RMSE occurs when REHE corrects negative values in HE estimates. We counted the proportion of simulation replicates where HE gave negative estimates (before zero truncation) for each setting, and observed a proportion as high as 45% ($n$ =3,000, $D_1$ under setting 2, $(\sigma_0^2,\ \sigma_1^2) \in \{(0.01, 0.1),\ (0.1, 0.01)\}$). Even at the sample size $n$ =12,000, HE still produces negative estimates in 30% of the replicates. Even with less extreme variance component values $(\sigma_0^2,\ \sigma_1^2) \in \{(0.04, 0.1),\ (0.1, 0.04)\}$, we still observed 15% of the HE estimates being negative at the sample size $n$ =3,000. In all of these cases, we obtained substantially lower RMSE by REHE than by HE (Figure S5-e, S5-f).

Confidence intervals have close to nominal coverage for all methods under $(\sigma_0^2, \sigma_1^2) = (0.1, 0.1)$, but show more volatility when the true variance component values are unequal (Figure S6). Taking into consideration the Monte Carlo error of 0.03 based on 200 replicates, REHE quantile-type confidence intervals always have acceptable coverage around 0.95. The coverage of REHE Wald-type confidence intervals is generally close to 0.95, but slightly off under some scenarios (Figure S6-c, S6-f, S6-i). This suggests that REHE quantile-type interval is more robust than Wald-type interval. REHE confidence intervals constructed from sparsified matrices have similar patterns of coverage. We observed in the main paper and Supplementary Note 2.1 that the coverage of REML and sREML is poor in some scenarios. Although the coverage improves quickly with increasing sample size in some cases, the under-coverage issue can persist even with large sample sizes (Figure S6-c, S6-e, S6-h). With $(\sigma_0^2, \sigma_1^2) = (0.01, 0.1)$ and kinship matrix structure under setting 2 or 3, the coverage of REML/sREML is lower than 0.7 even at the sample size $n = 12,000$ (Figure S6-e, S6-h). In addition, REML may fail to produce a confidence interval due to computational issues, which happens when any variance component estimate reaches zero during the iterative updates. Under setting 2, with $\sigma_0^2 = 0.1$,$\sigma_1^2 = 0.01$, $n$=3,000, this issue occurs in more than 30% of the simulation replicates. Even at a sample size of $n$ =12,000 we still observed as high as 22% of replicates having problem with REML inference. On the other hand, REHE confidence intervals does not suffer from this issue and can always be obtained.

Figure S7 illustrates the distribution of confidence interval half width. Overall REHE is comparable to REML. REML generally has narrower confidence intervals when true values of variance components are not very different (Figure S7-a, S7-d, S7-g), when they do differ REHE intervals are narrower (Figure S7-c, S7-f, S7-i). As we have seen scenarios with poor coverage for REML and sREML confidence intervals (Figure S6-e, S6-h), corresponding interval width has suspiciously skewed distributions (Figure S7-e, S7-h).

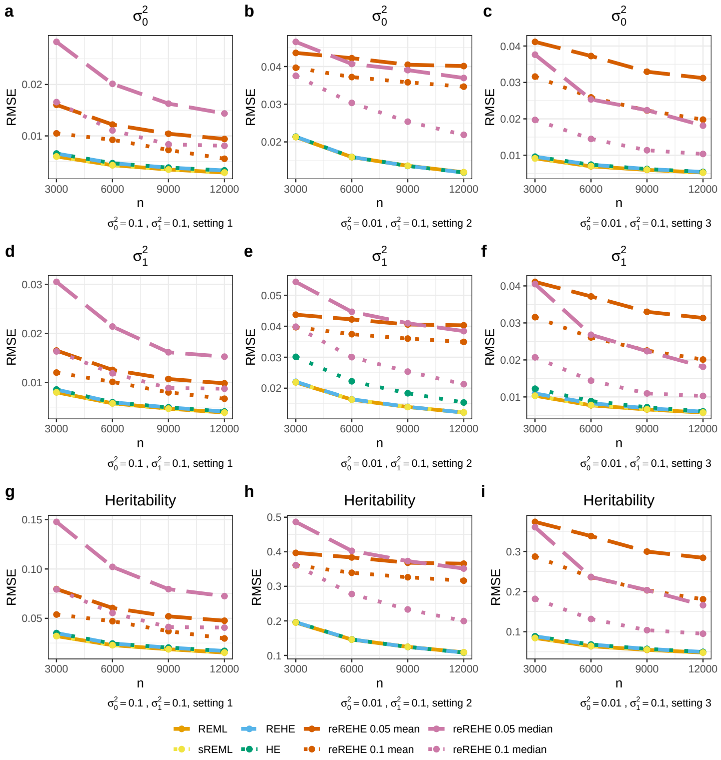


Figure S5: Results of simulation study on heritability analysis. RMSE of estimates for $\sigma_0^2$, $\sigma_1^2$ and heritability, based on REML, REML with matrix sparsification (sREML), REHE, HE, and reREHE with different subsampling rates (0.05/0.1) and different summary functions (mean/median). Parameter true values for $\sigma_0^2$ and $\sigma_1^2$, and kinship matrix structure setting under each scenario are indicated on the bottom-left corner of each sub-plot. The title of each sub-plot indicates the parameter of interest.

### 2.4 Simulation Results on Network-Based Pathway Enrichment Analysis

We conducted additional simulation studies with network-based pathway enrichment analysis [3] to benchmark the performance of REHE and reREHE against REML. We designed the simulation study based on the breast cancer data and the dysregulation framework used in [4]. The breast cancer data set has 403 samples in the ER positive group, 117 samples in the ER negative group, with a total of 2,598 gene entries spanning 100 gene pathways. To minimize the difference in gene-gene correlations between the two groups, we randomly split the total number of 520 samples into one subset with 403 samples and another with 117 samples. The first subset of samples was used to estimate an influence matrix $\Lambda_1$ for the ER positive group, and the second an influence matrix $\Lambda_2$ for the ER negative group [3]. Next, we generated observations according to the linear mixed model (S2), where $T = 2$, $n_1 = 403$, $n_2 = 117$, and $p$ =2,598. We set the mean signal vector to be zero for the ER positive group: $\mu_{1,j} = 0$ for $j = 1, \dots, p$. For the ER negative group, we set $\mu_{2,j} = \Delta\mu \in \{0.1, 0.2, 0.3\}$ when $j \in S$ for an index set $S$, otherwise $\mu_{2,j} = 0$. This index set $S$ includes genes selected in the betweeness-type dysregulation design [4]. The true variance component values were set to be $(\sigma_0^2,\ \sigma_1^2) = (0.1, 0.1),\ (0.5, 0.1),\ (0.1, 0.5),\ (0.1, 1),\ (1, 0.1)$. Based on the semi-synthetic data, we performed network-based pathway enrichment analysis to compare the power detecting differential pathway against the true power for each pathway, where true powers are computed according to [3]. We also evaluated the variance component estimates against the true values. Under each setting we ran 200 replicates. All methods were implemented using R package *netgsa* (*v3.1.0*, 3).

Computation with REML is time-demanding, which took around 1.5 hours for the analysis, compared to only 2 minutes by REHE and reREHE based analysis. In addition to computational gain, REHE provides more accurate variance component estimates than REML (Figure S8). When the noise variance component $\sigma_0^2 = 0.1$ and the random effect variance component $\sigma_1^2 = 1$, REML estimates are far from truth, but REHE and reREHE estimates are accurate. This suggests that in practice one needs to be cautious about REML estimates if the majority of the outcome variation is attributed to the network random effects. reREHE estimates are comparable to REHE in general, but less accurate when the noise variance component is larger than the random effect variance (Figure S8).

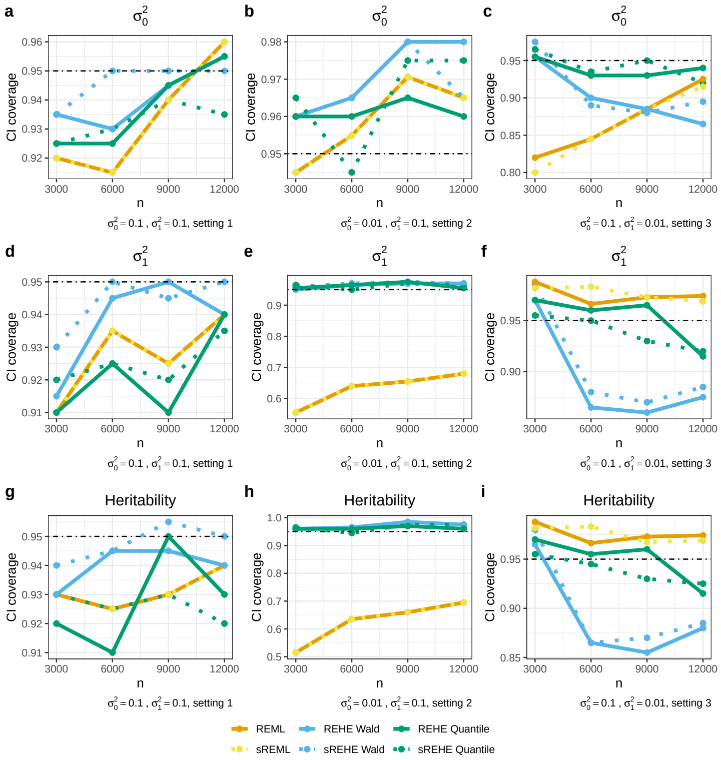


Figure S6: Results of simulation study on heritability analysis. Confidence interval coverage for $\sigma_0^2$, $\sigma_1^2$ and heritability with 200 replicates and expected Monte Carlo error of 0.03. We present results for confidence intervals constructed based on REML, REML with matrix sparsification (sREML), Wald-type intervals with REHE (REHE Wald), Wald-type intervals with REHE and matrix sparsification (sREHE Wald), quantile-type intervals with REHE (REHE Quantile), and quantile-type intervals with REHE and matrix sparsification (sREHE Quantile). Parameter true values for $\sigma_0^2$ and $\sigma_1^2$, and kinship matrix structure setting under each scenario are indicated on the bottom-left corner of each sub-plot. The title of each sub-plot indicates the parameter of interest.

Although REML provides less accurate variance component estimates than REHE, its empirical power for each pathway is closer to the true power in most settings, except when $(\sigma_0^2,\ \sigma_1^2) = (1, 0.1)$. In this case, the REHE approach has powers closer to the truth (Figure S9). reREHE empirical powers have similar patterns as those of REHE, but are slightly worse (Figure S9). In general the empirical powers of all three methods are close to the truth.

We noticed that even when REML fails to provide accurate variance component estimates, its empirical powers are still very close to the true powers. This indicates potential unidentifiability of the model parameters over the likelihood surface. This makes REML difficult to estimate the parameters, but it may still retain sensitivity of tests to group difference based on REML estimates. However, both REHE and reREHE can always yield good empirical powers and at the same time provide good estimates of variance components.

### References

[1] Matthew P. Conomos, Stephanie M. Gogarten, Lisa Brown, Han Chen, Ken Rice, Tamar Sofer, Timothy Thornton, and Chaoyu Yu. *GENESIS: GENetic EStimation and Inference in Structured samples (GENESIS): Statistical methods for analyzing genetic data from samples with population structure and/or relatedness*, 2019. R package version 2.14.3.

[2] Stephanie M Gogarten, Tamar Sofer, Han Chen, Chaoyu Yu, Jennifer A Brody, Timothy A Thornton, Kenneth M Rice, and Matthew P Conomos. Genetic association testing using the genesis r/bioconductor package. *Bioinformatics*, 35(24):5346–5348, 2019.

[3] Jing Ma, Ali Shojaie, and George Michailidis. Network-based pathway enrichment analysis with incomplete network information. *Bioinformatics*, 32(20):3165–3174, 2016.

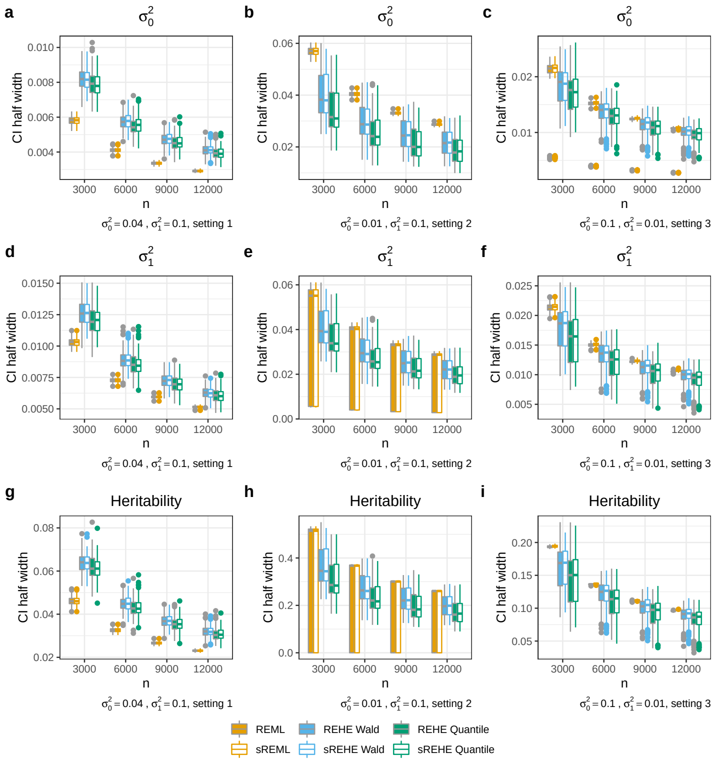


Figure S7: Results of simulation study on heritability analysis. Boxplots of confidence interval half width for $\sigma_0^2$, $\sigma_1^2$ and heritability. The boxplots follow standard Tukey representations, where center line represents median, box limits represent lower and upper quartiles, whiskers represent 1.5x interquartile range, and points represent outliers. We present results for confidence intervals constructed based on REML, REML with matrix sparsification (sREML), Wald-type intervals with REHE (REHE Wald), Wald-type intervals with REHE and matrix sparsification (sREHE Wald), quantile-type intervals with REHE (REHE Quantile), and quantile-type intervals with REHE and matrix sparsification (sREHE Quantile). Parameter true values for $\sigma_0^2$ and $\sigma_1^2$, and kinship matrix structure setting under each scenario are indicated on the bottom-left corner of each sub-plot. The title of each sub-plot indicates the parameter of interest.

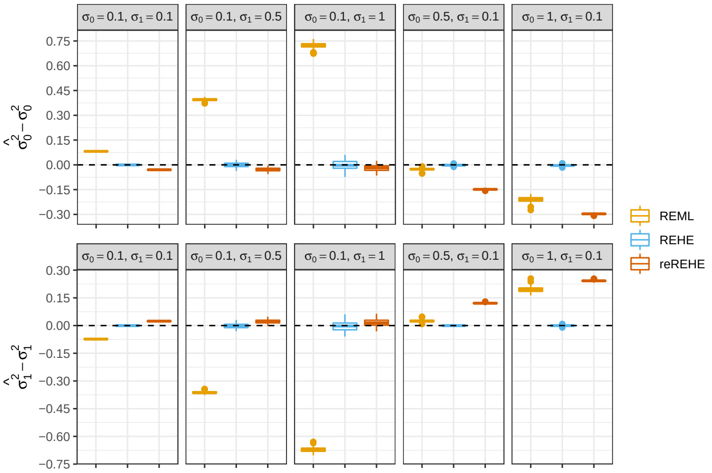


Figure S8: Results of simulation study on gene-network enrichment analysis. We present results for REML, REHE and reREHE, with boxplots of difference between the variance component estimates and the true values. The boxplots follow standard Tukey representations, where center line represents median, box limits represent lower and upper quartiles, whiskers represent 1.5x interquartile range, and points represent outliers. We present the results based on $\Delta\mu = 0.1$, and sub-plot titles indicate the parameter values of each setting. Results are based on 200 replicates.

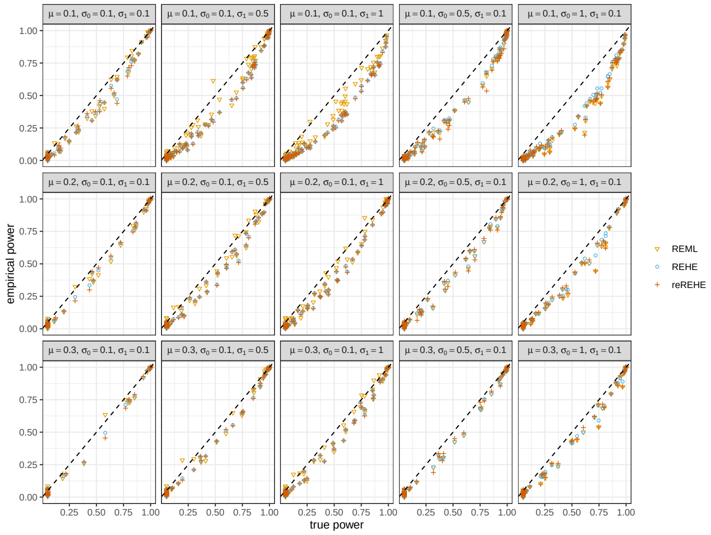

Figure S9: Simulation study results on gene-network enrichment analysis. We present empirical power against true power of detecting group difference for each pathway, for REML, REHE and reREHE approaches. Results are based on 200 replicates. The sub-plot titles indicate the parameter values of each setting.

[4] Jing Ma, Ali Shojaie, and George Michailidis. A comparative study of topology-based pathway enrichment analysis methods. *BMC bioinformatics*, 20(1):546, 2019.

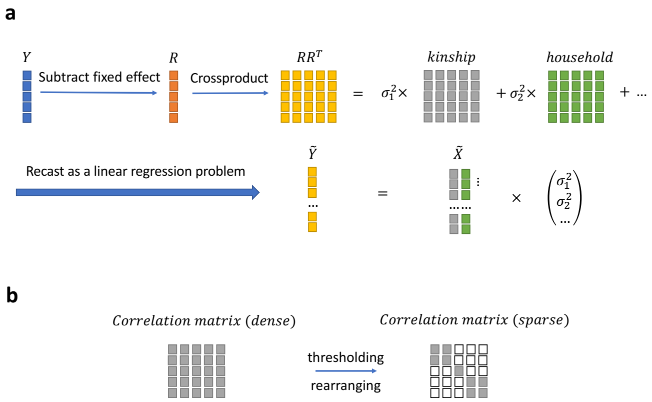


Figure S10: Illustration of HE/REHE workflow and correlation matrix sparsification. **a** - HE/REHE workflow. $Y$ is the outcome vector and $R$ is the residual vector after subtracting fixed effects from $Y$. Estimating equations are formed by decomposing the total variation $RR^\top$ into different sources (e.g. kinship, household relatedness) with corresponding variance components $\sigma_k^2$'s. After vectorizing the total variation $RR^\top$ as $\tilde{Y}$ and the relatedness matrices as $\tilde{X}$, variance components are estimated using ordinary least squares/non-negative least squares. **b** - Illustration of correlation matrix sparsification. Grey area represents non-zero entries, and white area represents zero entries.
